# Chronic trazodone treatment consolidates sleep, improves memory, and reduces amyloid pathology in a mouse model of Alzheimer’s disease

**DOI:** 10.64898/2026.08.20.746036

**Authors:** Mayuko Arai, Jefferey Yue, Emad Shams, Cody J Stevens, Hillary Han, Robert Gibson, Taha Yildirim, Howard H Feldman, Cheryl L Wellington, Brianne A Kent

**Affiliations:** Department of Psychology, Simon Fraser University, Burnaby, Canada; Institute for Neuroscience and Neurotechnology, Burnaby, Canada; Department of Molecular Biology and Biochemistry, Simon Fraser University; Department of Neurosciences, University of California, San Diego, La Jolla, USA; Department of Pathology and Laboratory Medicine, University of British Columbia, Vancouver, British Columbia, Canada

**Author notes:** Corresponding Author: Brianne A Kent, Department of Psychology, Simon Fraser University, Burnaby, Canada.

**Keywords:** sleep, Alzheimer’s disease, transgenic mouse model, amyloid, trazodone, memory, drug repurposing

## Abstract

Sleep disturbance in Alzheimer’s disease (AD), particularly the reduction of slow wave sleep (SWS), has been proposed as a novel therapeutic target, with disease-modifying potential. Trazodone, an antidepressant with robust SWS-promoting properties, is currently the most prescribed sleep-promoting medication in the United States. Here, we demonstrate that chronic trazodone administration consolidates sleep in the APP^NL-F^ knock-in mouse model of AD, increasing NREM sleep duration and slow wave power during the rest phase while promoting wake during the active phase. These sleep consolidating effects were accompanied by lower regional glial activation and amyloid burden, particularly in male mice. Most notably, hippocampal amyloid plaque burden was 45% lower in mice treated from 14 to 16 months of age than in vehicle-treated controls. Chronic trazodone treatment was also associated with better short-term and long-term recognition memory. Together, these findings support the potential of repurposing trazodone as a well-tolerated, disease-modifying therapeutic for AD, capable of enhancing sleep quality, improving cognition, and lowering AD-relevant neuropathology.

## Introduction

Alzheimer’s disease (AD) is the most common cause of dementia, defined by neuropathological hallmarks of extracellular deposits of β-amyloid (Aβ), neurofibrillary tangles composed of hyperphosphorylated tau, and neurodegeneration, which can start to develop decades before the onset of clinical symptoms^1^. The preclinical and prodromal stages of AD provide a critical window for early intervention to prevent the onset of cognitive decline and dementia.

Sleep is an emerging treatment frontier in AD^2^. Most individuals with AD experience some degree of sleep disturbance, manifesting as fragmented nighttime sleep, excessive daytime napping, and reductions in both slow-wave sleep (SWS) and rapid-eye-movement (REM) sleep^2^. Sleep has been shown to directly regulate the hallmark neuropathology of AD, both Aβ and tau in mouse models and humans^3–7^. A meta-analysis estimated that approximately 15% of AD cases in the population could be delayed or prevented by interventions targeting sleep^8^.

A promising therapeutic sleep target for AD is SWS, which dominates the N3 stage of non-rapid eye movement (NREM). Slow-wave activity during SWS is thought to reduce neuronal activity–dependent Aβ release ^9^, promote clearance of Aβ via CSF flow ^5,10–12^, and may mediate Aβ-associated impairment of hippocampus-dependent memory consolidation^13^. Individuals with AD and mouse models of AD commonly exhibit reduced SWS^14–16^. In older adults without dementia, low SWS is associated with AD-related biomarkers (cortical amyloid burden^13,17^, CSF Aβ42^18^, and cortical tau accumulation^19^) and increased dementia risk^20^. In mouse models, restoring SWS using optogenetic stimulation^21^ or zolpidem^22^ reduced Aβ deposition, and enhancing SWS with lemborexant reduced tau pathology^23^. Together, these studies from mice and humans provide strong support for SWS as a promising therapeutic target in AD.

Trazodone is an atypical anti-depressant with multimodal mechanisms of action, however, approximately 85% of its prescribed uses are for off-label indications, including insomnia^24^. It is currently the most prescribed hypnotic medication in the United States^25^. At lower doses (25–100 mg), trazodone causes antagonism of serotonin 5-HT2A, H1 histamine, and α1 adrenergic receptors, producing hypnotic effects, whereas at higher doses (150–600 mg) trazodone blocks serotonin transporters (SERTs), causing antidepressant effects^26^. Importantly, trazodone increases SWS and, at the low-doses used for targeting sleep, is well-tolerated by elderly individuals^27–29^. This is unlike many other commonly used sleep-wake modulating drugs including benzodiazepines or zolpidem that have motor or cognitive side effects making them unsuitable for elderly individuals or those diagnosed with neurodegenerative disorders^29–32^. Clinical research on trazodone in the context of AD is sparse but shows that trazodone improves sleep and may slow cognitive decline in individuals diagnosed with AD^27,32–35^.

Here, we show that chronic trazodone treatment has disease-modifying effects in a mouse model of AD, enhancing slow-wave sleep, reducing hippocampal Aβ pathology, and improving memory. Sixty days of daily trazodone treatment promoted sleep consolidation in both female and male mice, characterized by increased NREM sleep with higher slow-wave power during the rest phase and increased wakefulness during the active phase. Histological analyses revealed lower Aβ plaque development in both sexes and reduced regional glial activation in male mice. Trazodone may also attenuate microglial responses surrounding smaller fibrillar-cored plaques. In mice treated for 130 days, trazodone was associated with improved short-term and long-term recognition memory. Treatment was delivered using our previously validated voluntary oral administration protocol^36^.

## Results

### Chronic trazodone extends rest-phase NREM duration and consolidates sleep

We first examined the effects of trazodone on the duration of each vigilance state (NREM, REM, and wake). Animals were recorded with EEG/EMG while receiving trazodone (60 mg/kg) (treatment) or vehicle (control) daily for 60 days, beginning at 9 or 14 months of age (Fig. 1). During the light period (rest phase, ZT0–12), both the 9-month and 14-month cohorts treated with trazodone showed increased NREM duration (Fig. 2a, f), accompanied by reduced REM (Fig. 2b, g) compared to baseline. During the dark period (active phase, ZT12–23, excluding the last hour of the dark period, when dosing occurred), trazodone-treated animals exhibited a strong wake-promoting effect (Fig. 2c, h), along with a corresponding decrease in NREM sleep (Fig. 2a, f).

**Fig 1:**
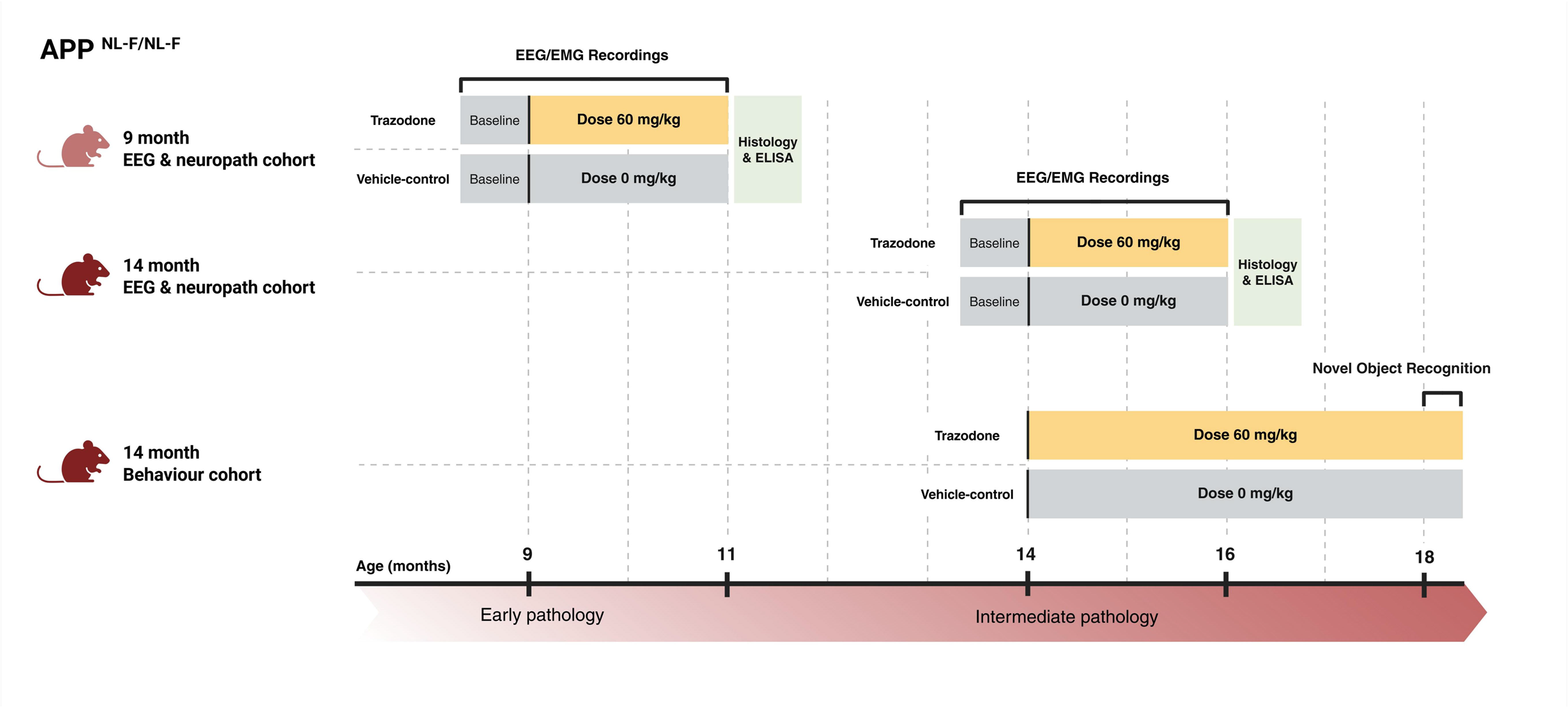
Timeline. Three cohorts of homozygous APP^NL-F^ mice were used. All cohorts were divided into two groups: a trazodone treatment group and a vehicle control group. The EEG/EMG and neuropathology cohorts began treatment at 9 months or 14 months of age, representing early and intermediate stages of amyloid pathology, respectively. Following the baseline EEG recording day when all mice received palatable food without trazodone, the treatment group received palatable food mixed with trazodone (60 mg/kg), and the control group received palatable food mixed with distilled water. During the 60 days of daily treatment, EEG was recorded from all animals for 2-day sessions (1 day habituation, 1 day recording) every 7–10 days. After the 60-day treatment, all animals were euthanized for histology and ELISA analyses. A separate behavioral cohort began daily trazodone or vehicle treatment at 14 months of age and underwent novel object recognition testing after approximately 130 days of treatment, at 18 months of age.

**Fig 2:**
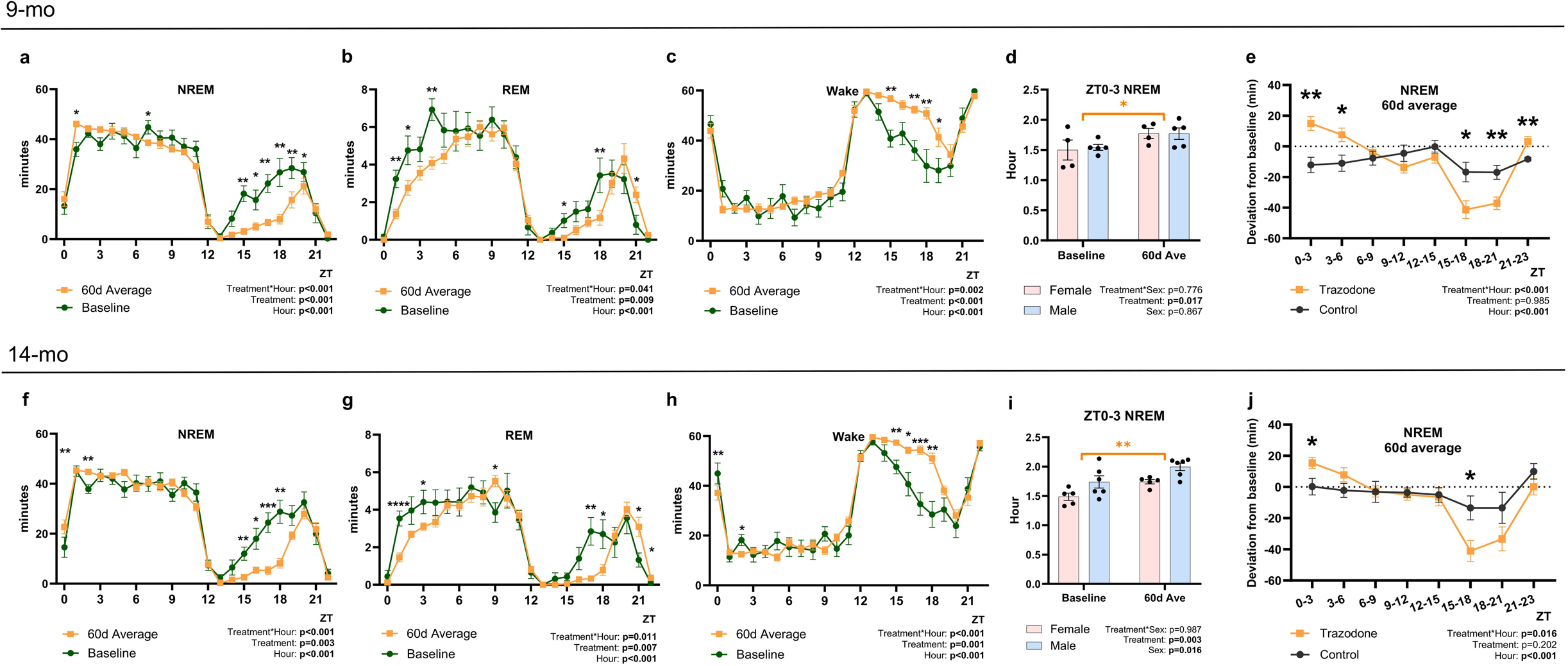
Trazodone increases NREM duration immediately after dosing and consolidates sleep. Effects of chronic trazodone treatment on vigilance state durations in APP^NL-F^ mice treated daily from 9 to 11 months of age (a–e) and from 14 to 16 months of age (f–j). **(a – c & f – h)** Duration of each vigilance state per hour, Baseline vs 60-day average. In both the 9-month (a–c) and 14-month (f–h) cohorts, trazodone was administered 30 min before lights-on at ZT0. In both cohorts, NREM duration increased at the start of the 12-h light phase at the expense of REM, while wake duration increased during the 12-h dark phase, when mice are normally more active. **(d & i)** Comparison across sexes of NREM at baseline versus the 60-day average. Analysis focused on ZT0-3, where significant NREM enhancement was observed in both cohorts. Sex differences were present at baseline, but the effects of trazodone did not differ between sexes. **(e & j)** NREM duration deviation from baseline every 3 h, Control vs Trazodone (60-day average). In both cohorts, a significant Treatment × Hour interaction was observed. Post hoc analysis showed that trazodone-treated animals had greater NREM enhancement at the start of the 12-h light phase and reduced NREM during the dark phase. Data (mean ± SEM) were analyzed by two-way ANOVA followed by Tukey’s multiple comparisons test, or mixed-effects models when there were missing values, followed by Šidák’s multiple comparisons tests. * p<0.05, ** p<0.01, *** p<0.001.

To closely assess the NREM-promoting effect, we analyzed NREM duration during the first three hours after dosing (ZT0–3), separated by sex (Fig. 2d, i). In the 9-month cohort, a two-way ANOVA (treatment × sex) revealed a significant main effect of treatment, showing higher NREM duration during the 60-day treatment, with no significant effect of sex (Fig. 2d). In the 14-month cohort, both treatment and sex showed significant main effects, indicating that trazodone increased NREM duration and that males spent longer time in NREM regardless of treatment (Fig. 2i).

Finally, we examined the deviation in NREM duration from baseline between trazodone- and vehicle-treated groups, averaged across 3-hour intervals over the 60-day period (Fig. 2e, j). In both the 9-month (Fig. 2e) and 14-month (Fig. 2j) cohorts, a significant treatment × time interaction was observed (p < 0.001). Post hoc analysis showed that trazodone increased NREM sleep during the early light phase and decreased NREM during the dark phase in both cohorts, compared to the vehicle control (Fig. 2e, j).

Together, these results suggest that trazodone treatment consolidates sleep, promoting NREM sleep immediately after administration at the start of the rest (light) phase and enhancing wakefulness during the active (dark) phase. Trazodone’s effects on vigilance state durations were consistent for both sexes.

### Trazodone increases EEG slow-wave power

In addition to observing the increase in NREM duration immediately after trazodone dosing, we, next, analyzed changes in EEG power in the first 3 hours after dosing (ZT0-3). In both the 9-month and 14-month cohorts, normalized NREM EEG power spectra showed a significant increase in slow oscillation (SO, 0.5–1 Hz) and delta (1–4 Hz) power, accompanied by a reduction in theta power, during the 60 days of dosing compared to baseline (Fig. 3a, f). Similarly, in both cohorts, REM EEG power spectra shifted toward lower frequencies, with increased delta power and reduced alpha power over the 60-day treatment period (Fig. 3b, g).

**Fig 3:**
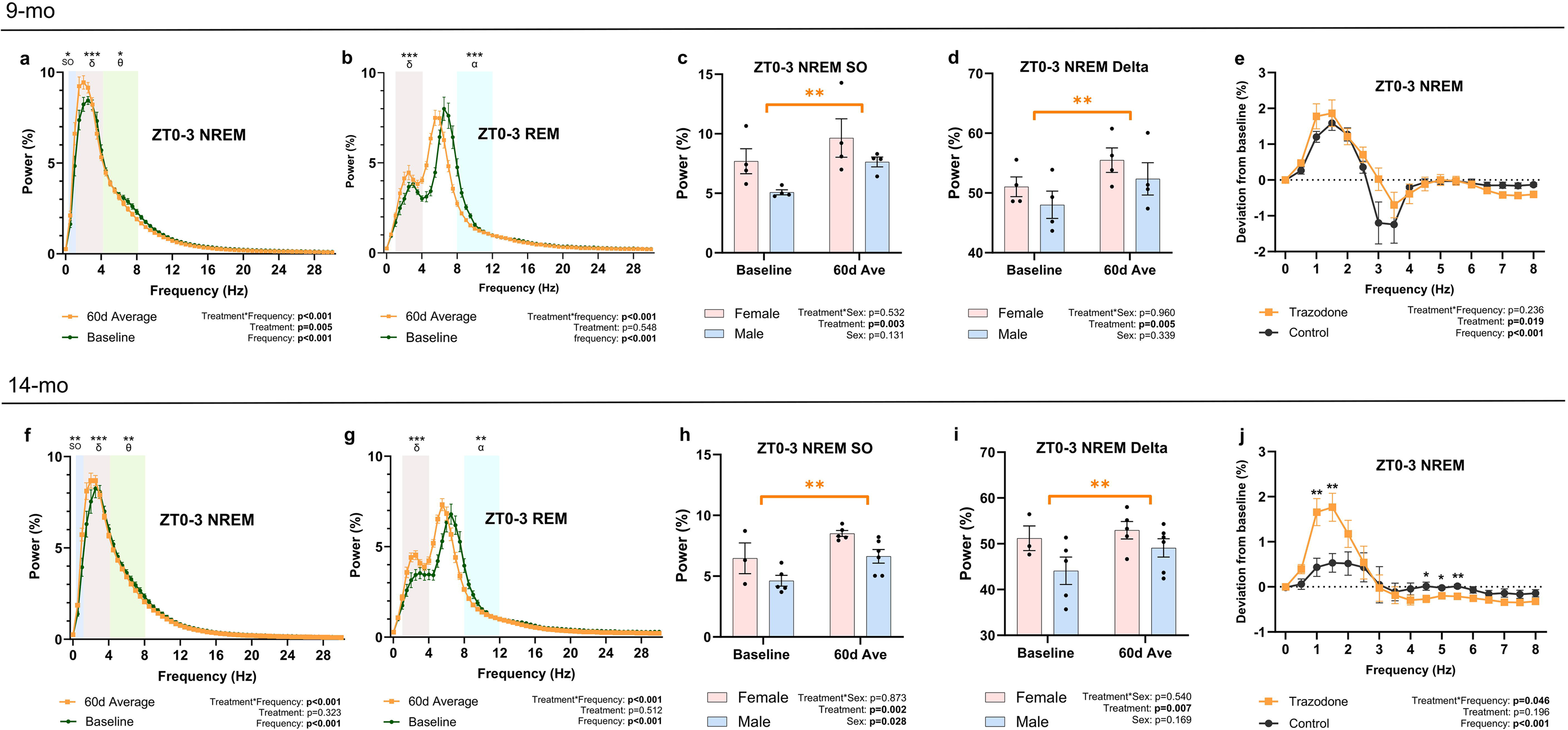
Trazodone shifts the power spectra immediately after dosing. Effects of chronic trazodone treatment on frontal EEG power during ZT0-3 in APP^NL-F^ mice treated daily from 9 to 11 months of age (a–e) and from 14 to 16 months of age (f– j). **(a, f)** ZT0-3 NREM normalized power spectra of trazodone-treated animals at baseline and averaged across the 60 days of treatment. During treatment, SO and delta power increased, accompanied by a reduction in theta power in both cohorts. **(b, g)** ZT0-3 REM normalized power spectra of trazodone-treated animals at baseline and averaged across the 60 days of treatment, showing a shift of the power spectrum toward lower frequencies. **(c, d, h, i)** Comparison across sexes of ZT0-3 NREM SO and delta power, at baseline and averaged across the 60 days of treatment. Sex differences were observed at baseline, but the effects of trazodone did not differ between sexes. **(e & j)** ZT0–3 NREM power deviation from baseline in vehicle-control versus trazodone groups (60-day average), showing that trazodone-treated animals from 14 to 16 months of age exhibited greater slow-wave power, particularly between 1 and 1.5 Hz. Data (mean ± SEM) were analyzed by two-way ANOVA followed by Tukey’s multiple comparisons test, or mixed-effects models when there were missing values, followed by Šidák’s multiple comparisons tests. * p<0.05, ** p<0.01, *** p<0.001.

Given the prominent changes in SO and delta power, we analyzed these bands separately by sex. When averaged across ZT0–3, a two-way ANOVA (treatment × sex) showed a significant treatment effect, with both cohorts exhibiting higher SO and delta power during the 60-day treatment than baseline (Fig. 3c, d, h, i). No significant sex differences were observed in the 9-month cohort, although females showed a higher mean SO and delta power than males (Fig. 3c, d). In the 14-month cohort, females exhibited significantly higher SO power than males regardless of treatment (Fig. 3h).

To further examine frequency-specific changes, the deviation in power from baseline was averaged across ZT0–3 and compared between trazodone- and vehicle-treated control groups. In the 9-month cohort (Fig. 3e), although a significant main effect of treatment was observed (p = 0.019), no specific frequency bin showed a clear difference. In contrast, in the 14-month cohort (Fig. 3j), a significant treatment × frequency interaction was found (p=0.046). Post hoc analysis revealed that trazodone-treated mice in 14-month cohort showed greater slow-wave power enhancement, particularly between 1 and 1.5 Hz, accompanied by reduced power in the 4.5–5.5 Hz range.

We next examined how these effects on slow-wave power changed throughout the day (excluding the final dark-phase hour used for dosing). When the 60-day average power was compared to baseline, SO (0.5-1Hz) power during NREM sleep was significantly increased during the 12-h light period in both the 9-month and 14-month cohorts (Extended Data Fig. 1a, e). The effect was most pronounced at the start of the light period. A similar pattern was observed for delta (1–4 Hz) power in both cohorts, although the difference was only statistically significant in the 14-month group (Extended Data Fig. 1b, f). When the 60-day average power was compared to the vehicle-treated group, a similar increase in SO and delta power was observed in both cohorts at the beginning of the 12-h light period (Extended Data Fig. 1c, d, g, h).

Next, we analyzed these bands separately by sex. When SO and delta power averaged across the 23-h period were compared with baseline, a two-way ANOVA (treatment × sex) revealed significantly elevated SO power in both cohorts. Females showed higher SO power than males in both cohorts, although this difference reached statistical significance in only in the 14-month cohort (Extended Data Fig. 2a, d). A similar effect was seen for delta power (Extended Data Fig. 2b, e).

Finally, the deviation in power from baseline was averaged across 23 hours and compared between trazodone-treated and vehicle-treated groups. The 23-h average showed modest changes in SO and delta power, although were not statistically significant in the 9-month (Extended Data Fig. 2c) or 14-month (Extended Data Fig. 2f) cohort. Taken together, trazodone increased SO and delta power during NREM sleep at the start of the light period in both sexes.

### Chronic trazodone reduces amyloid-plaque accumulation

We assessed amyloid burden following 60 days of trazodone treatment by histological and biochemical analyses. In the 9-month cohort (Fig. 4a, c, e), immunohistochemistry revealed a main treatment effect in the entorhinal cortex (p=0.004, F(1, 12) = 12.58), indicating 32% lower amyloid plaque burden. In the 14-month cohort (Fig. 4i, k, m), trazodone treatment was associated with 45% lower amyloid plaque in the hippocampus (p=0.011, F(1, 15) = 8.497). We then assessed the effects of trazodone in the deposition of fibrillar-cored amyloid plaques by staining with fluorogenic dye X-34 (Extended Data Fig. 3) and Thioflavin-S (Fig. 4 b, d, f, j, l, n). Two-way ANOVA analysis (treatment x sex) revealed a trend toward interaction effect on X-34 labeled plaques in the entorhinal cortex of the 14-month cohort after trazodone treatment (p=0.068), with post hoc analysis suggesting reductions in the male cohort (p=0.037, t(14) = 2.659; Extended Data Fig. 3f). For Thioflavin-S assessment, no clear treatment effects were shown in the 9-month cohort (Fig. 4b, d, f); however, a consistent downtrend was found across all regions in the 14-month cohort (Fig. 4j, l, n), with the temporal cortex showing a statistically significant reduction between the vehicle and trazodone treated groups (p=0.033, F(1, 15) = 5.505).

**Fig 4:**
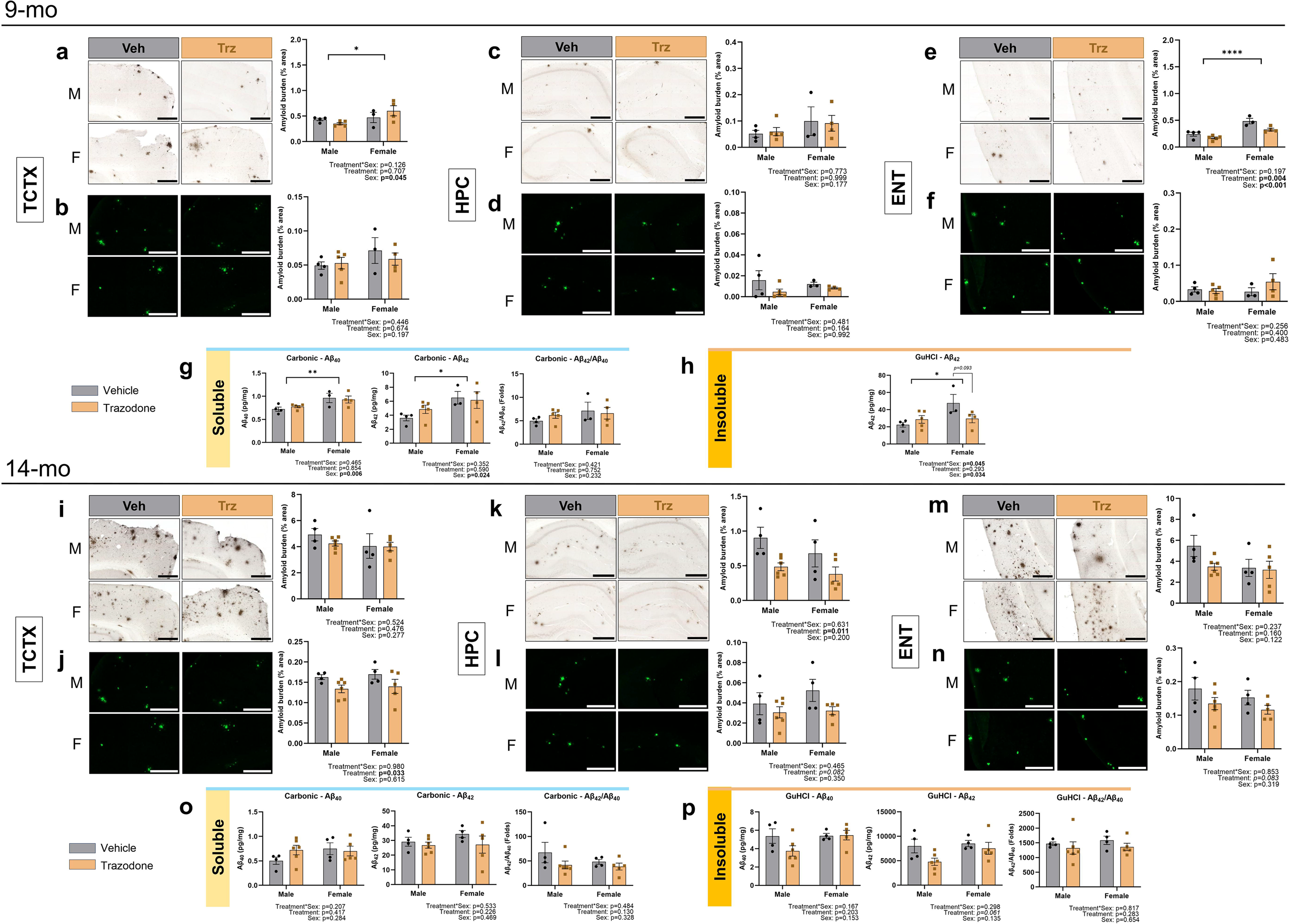
Trazodone reduces amyloid aggregate burden in the brain. Effects of chronic trazodone treatment on amyloid burden in APP^NL-F^ mice treated daily from 9 to 11 months of age (a–h) and from 14 to 16 months of age (i–p). Representative histological images showing reduction of amyloid plaque deposition in the brain. Antibody-labeled amyloid plaques in the **(a, i)** temporal cortex, **(c, k)** hippocampus, and **(e, m)** entorhinal cortex are shown. Thioflavin-S-labeled amyloid plaques in the **(b, j)** temporal cortex, **(d, l)** hippocampus, and **(f, n)** entorhinal cortex. The percentage of plaque area within each region are reported in the bar graphs. **(g, h, o, p)** Biochemical measurement of soluble and insoluble Aβ in brain lysate. ELISA assays quantifying **(g,o)** soluble and (**h,p)** insoluble Aβ42, Aβ40, and the corresponding Aβ42: Aβ40ratio. Each data point represents one sample. Data (mean ± SEM) were analyzed by two-way ANOVA followed by Bonferroni multiple comparison post-hoc test. * p<0.05, ** p<0.01. Scale bar for IHC = 500 µm. Scale bar for ThioS = 200 µm.

We also quantified soluble and insoluble Aβ by analyzing carbonic and GuHCl fractions of brain lysate by ELISA, respectively. An interaction effect (treatment x sex) on GuHCl-soluble Aβ42 was found in the 9-month cohort (p=0.045, F(1, 12) = 4.996), with the trazodone-treated female mice showing a downtrend (Fig. 4h). The concentration of GuHCl-soluble Aβ40 did not reach detection sensitivity and was therefore excluded from analyses in the 9-month cohort. No treatment effects were found in the carbonic-soluble Aβ (Fig. 4g,o). In the 14-month cohort, a downtrend in treatment effect was found in the GuHCl-soluble Aβ42 (p=0.061; Fig. 4p). We also assessed the Aβ42:Aβ40 ratio in all fractions which was thought to precede amyloid plaque formation.^37,38^ While males and females showed reductions in the Aβ42:Aβ40 ratio (Fig. 4o,p), none were statistically significant. An exploratory correlational analysis between neuropathology and NREM power revealed several negative associations between NREM SO and delta power and levels of Aβ40, Aβ42 (Extended Data Fig. 4a, c, d), and amyloid plaque burden (Extended Data Fig. 4h), such that the higher levels of slow-wave power across the 60 days was associated with lower pathology; however, these analyses were notably underpowered. Collectively, these findings suggest that trazodone attenuates the accumulation and aggregation of amyloid in differential patterns amid different stages of AD progression.

### Trazodone reduces regional glial activation in male mice

We next assessed the effect of chronic trazodone treatment on neuroinflammation by examining microglial activation in the APP^NL-F^ mice (Fig. 5). We found an interaction effect (treatment x sex) in Iba1-immunoreactivity in the entorhinal cortex of the 9-month cohort (p=0.013, F(1, 12) = 8.398; Fig. 5c), with post-hoc analysis indicating lower microglial activity in the male mice (p=0.009, t(12) = 3.476). In the 14-month cohort, all investigated regions showed downtrends in microgliosis in male mice, with the hippocampus showing a statistically significant difference (p=0.031, t(14) = 2.763; Fig. 5h). Immunofluorescence analysis of GFAP-immunoreactivity indicative of astroglial activation revealed a treatment effect in the entorhinal cortex of the 14-month cohort (p=0.031, F(1, 14) = 5.780) (Fig. 5l). The same region also showed an interaction trend, with the male mice showing lower astroglial activation when treated with trazodone compared to vehicle treatment (p=0.010, t(14) = 3.313), suggesting chronic trazodone treatment exerts anti-inflammatory effect in the brain.^42^

**Fig 5:**
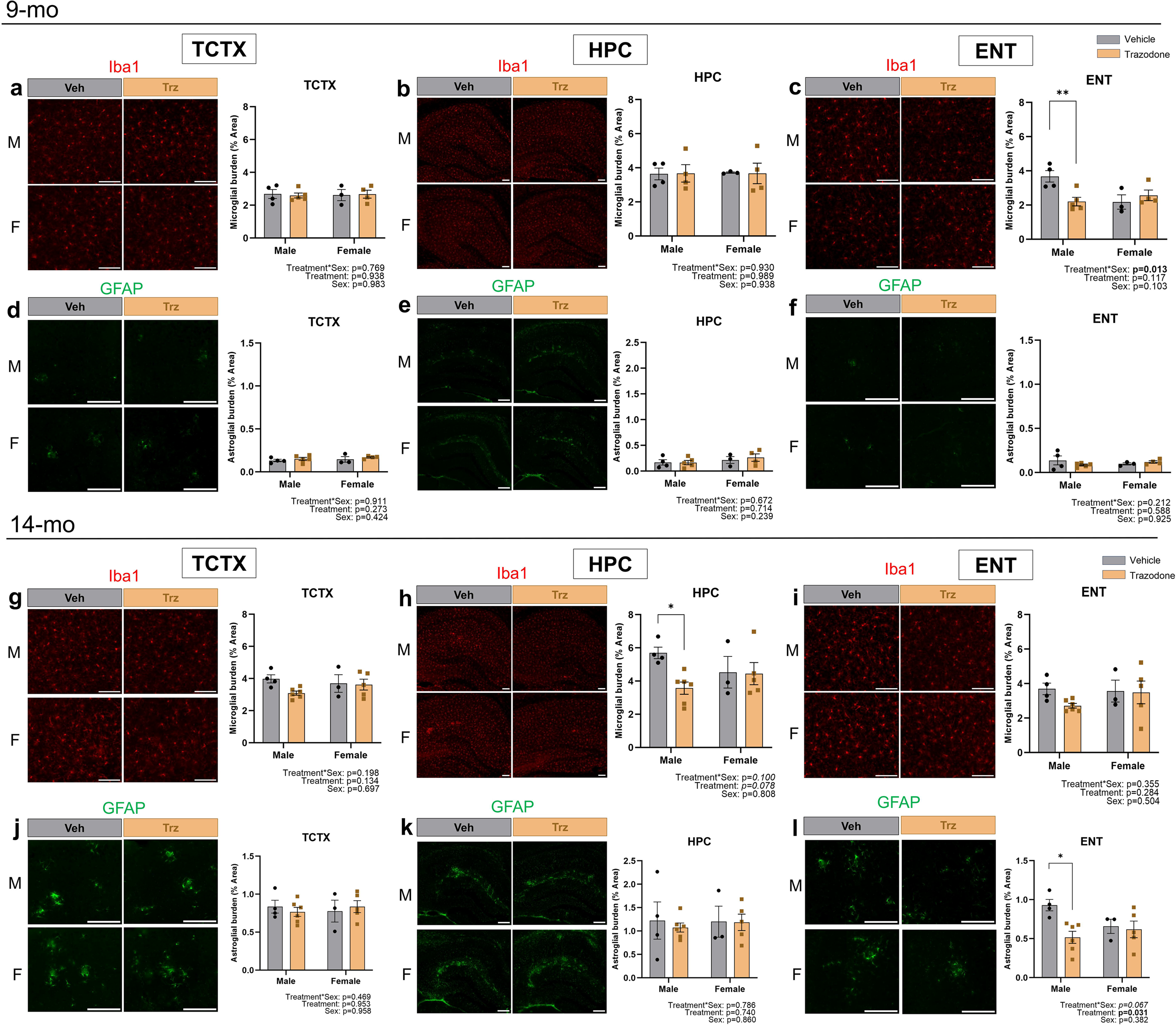
Trazodone exerts age-dependent dampening of glial activation. Effects of chronic trazodone treatment on microglial and astroglial activation in APP^NL-F^ mice treated daily from 9 to 11 months of age (a–f) and from 14 to 16 months of age (g– l). Representative immunofluorescence images showing reduced microglial activation in male mice after vehicle or trazodone treatment. Iba1 immunoreactivity in (**a, g**) temporal cortex, (**b, h**) hippocampus and (**c, i)** entorhinal cortex. Quantification of the percentage of Iba1 immunoreactive area in each region is reported in the bar graph. Representative immunofluorescence images showing reduced astroglial activation after trazodone treatment. GFAP immunoreactivity in **(d, j)** temporal cortex, **(e, k**) hippocampus, and **(f, l)** entorhinal cortex are shown. Quantification of the percentage of GFAP immunoreactive area in each region is reported in the bar graph. Each data point represents one sample. Data (mean ± SEM) were analyzed by two-way ANOVA followed by Bonferroni multiple comparison post-hoc test. * p<0.05, ** p<0.01. Scale bar = 200 µm.

### Trazodone dampens congophilic plaque-associated microglial activation in male mice

Given that microglial activation is significantly amplified while interacting with congophilic amyloid plaques,^39^ we further examined congophilic amyloid plaques-associated microglia by Iba1-immunoreactivity and X-34 staining to better understand whether trazodone affects microglia that are engaging mature amyloid aggregates (Fig. 6). In the 9-month cohort, we found an interaction effect on Iba1-immunoreactive area proximal to X-34 labeled plaques at the entorhinal cortex (p=0.021, F(1, 12) = 7.106; Fig. 6c), with post-hoc analysis suggesting significant decreases in male mice after trazodone treatment (p=0.029, t(12) = 2.849). In the 14-month cohort, two-way ANOVA revealed interaction effects (treatment x sex) at the hippocampus (p=0.038, F(1, 14) = 5.239; Fig. 6e) and entorhinal cortices (p=0.034, F(1, 14) = 5.530; Fig. 6f), with the male mice treated with trazodone showing significantly lower Iba1-immunoreactive area, than vehicle treated male mice (HPC p=0.009, t(14) = 3.387; ENT p=0.032, t(14) = 2.740).

**Fig 6:**
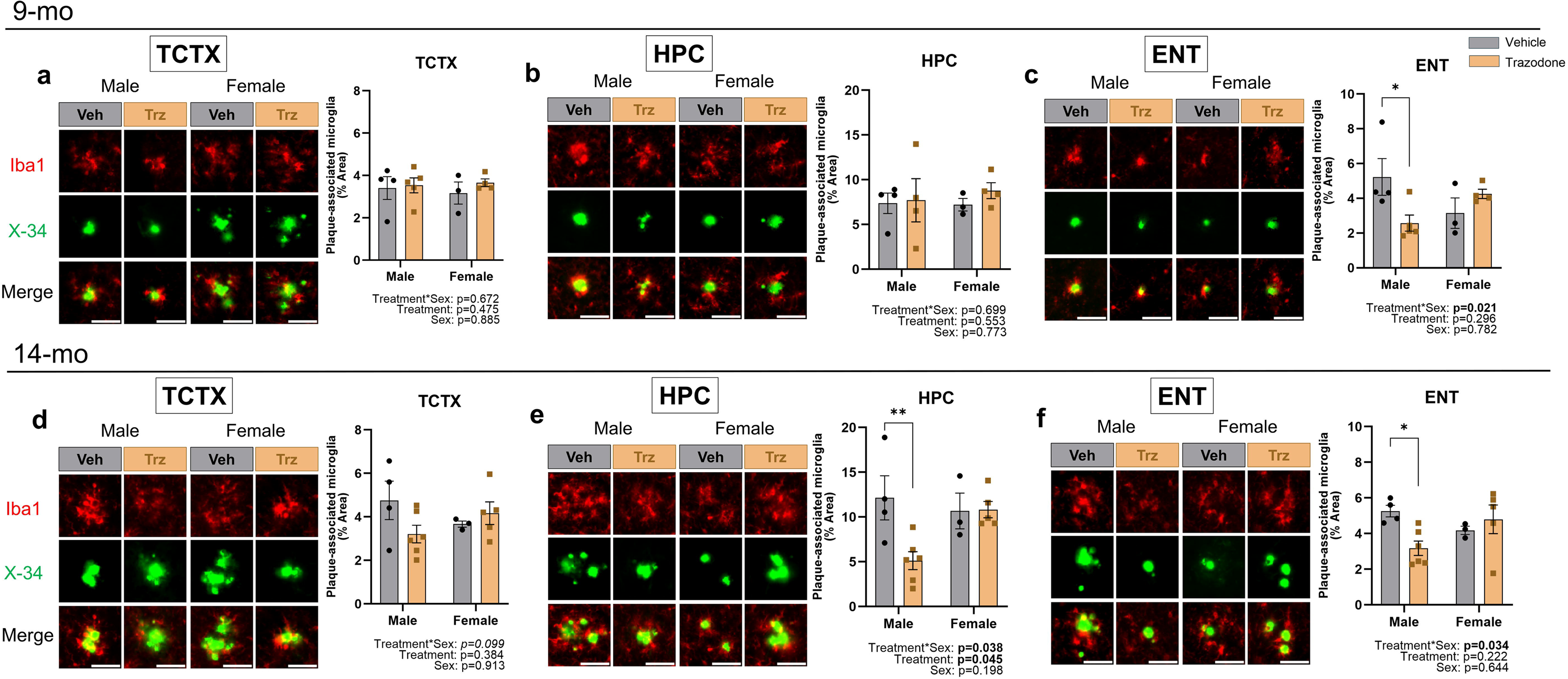
Trazodone lowers plaque-associated microglial activity in sex-dependent manner. Representative histological images showing sex-dependent reduction of microglial activity surrounding amyloid plaques in APP^NL-F^ mice treated daily from 9 to 11 months of age (a–c) and from 14 to 16 months of age (d–f). Antibody-labeled microglia and X-34 labeled fibrillary-cored amyloid plaques in the **(a,d)** temporal cortex, **(b,e)** hippocampus, and **(c,f)** entorhinal cortex. Quantification of Iba1-immunoreactive area within 50 µm from the parameter of X-34 labeled deposits within each region is reported in the bar graph. Each data point represents one sample. Data (mean ± SEM) were analyzed by two-way ANOVA followed by Bonferroni multiple comparison post-hoc test. * p<0.05, ** p<0.01. Scale bar = 50 µm.

### Trazodone lowers phosphorylated tau in the brain

We additionally measured endogenous murine total tau (t-tau) and phosphorylated tau (p-tau). Although the APP^NL-F^ mouse model does not develop hyperphosphorylated tau tangles, these measures may provide additional insight into early AD-related tau alterations. Unlike amyloid, the level of tau accumulation is not known to exhibit sex-dependent differences, so samples from both male and female animals were analyzed together. While no major changes on t-tau were found in either age group (Extended Data Fig. 5a–b, d–e), immunoblot analyses revealed significantly lower pS396/pS404 p-tau in the 9-month cohort treated with trazodone, compared to vehicle treated mice (p=0.021, t(14) = 2.593) but no significant treatment effects in the 14-month cohort (Extended Data Fig. 5c, f). Since this mouse model does not express tau mutants that exert aggregation propensity, we did not further assess regional tau burden by histological analysis. Our findings suggest that chronic trazodone treatment may reduce pathological phosphorylation of endogenous tau in the early stage of AD.

### Trazodone enhances short-term and long-term recognition memory

We assessed object recognition memory in a separate cohort of APP^NL-F^ that began daily treatment (60mg/kg trazodone or vehicle) at approximately 14 months of age and underwent novel object recognition testing after approximately 130 days of treatment at around 18 months of age. Recognition memory was evaluated after either a 1-h or 24-h delay, and discrimination indices (DIs) were calculated as the time spent exploring the novel object divided by the total object exploration time during the test phase.^40^

After a 1-h delay, two-way ANOVA revealed a significant main effect of treatment across sexes (F(1,24) =14.59, p<0.001; Fig. 7a). Similarly, after a 24-h delay, there was a significant main effect of treatment across sexes (F(1,25)=18.99, p<0.001; Fig. 7b). Together, these findings suggest that chronic trazodone treatment was associated with enhanced recognition memory at both short- and long-term retention intervals.

**Fig 7:**
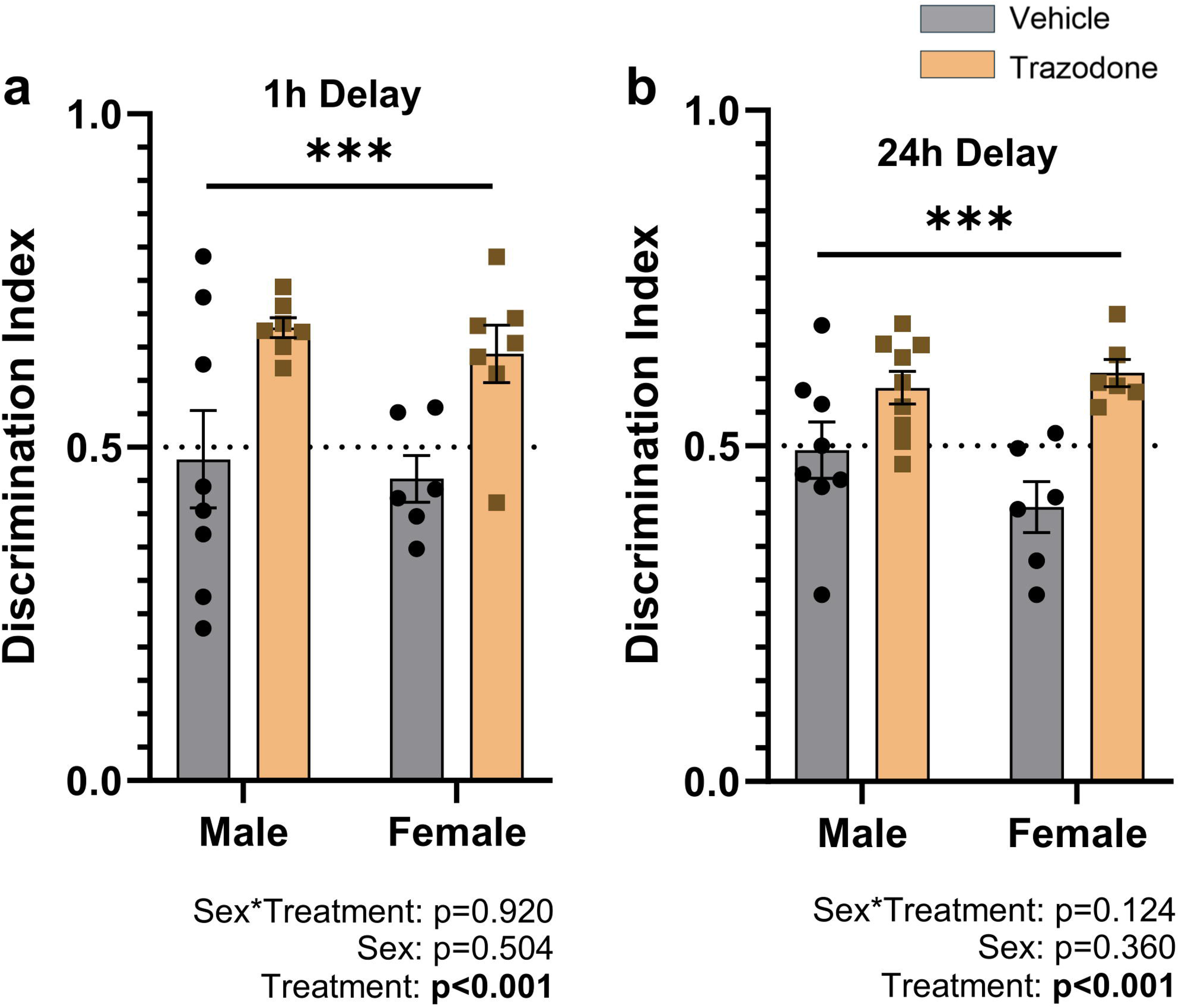
Chronic trazodone treatment is associated with improved short- and long-term recognition memory. Effects of chronic trazodone treatment on novel object recognition memory in APP^NL-F^ mice treated daily from 14 to 18 months of age. Testing was performed after approximately 130 days of treatment using delays of (a) 1 h and (b) 24 h. The discrimination index was calculated as the time spent exploring the novel object divided by the total time spent exploring both objects during the test phase. The dashed line at 0.5 represents chance-level performance (equal exploration of the novel and familiar objects); values above 0.5 indicate a preference for the novel object. Data (mean ± SEM) were analyzed by two-way ANOVA. *** p<0.001.

## Discussion

Trazodone is being explored as a repurposed therapeutic candidate for AD. Here, we provide novel evidence supporting trazodone’s potential as a multimodal, disease-modifying therapeutic for AD by consolidating sleep, improving cognition, dampening neuroinflammation, and lowering amyloidogenic pathology. Specifically, in the APP^NL-F^ knock-in mouse model of AD, chronic trazodone treatment increased NREM duration and NREM slow-wave power immediately after dosing, which had a consolidating effect, promoting wake during the active phase. The active-phase wake-promoting effect was an unexpected benefit of improved sleep quality, and one with clear clinical value. Accompanying the sleep enhancement, 60 days of trazodone treatment was also associated with lower AD neuropathology. Specifically, mice treated with trazodone exhibited lower regional microglial and astroglial activation, lower amyloid aggregates, and attenuated microglial activity associated with fibrillar-cored plaques, particularly in male mice. We also observed lower levels of phosphorylated tau at S396/pS404 – a p-tau species prominent in neurofibrillary tangles – in mice expressing early stage of amyloid pathology. In addition to the beneficial effects on sleep and neuropathology, trazodone-treated mice demonstrated superior short-term and long-term recognition memory, a cognitive domain compromised early in the course of AD.

These results add to the growing body of clinical and preclinical evidence supporting beneficial effects of trazodone in AD by providing the first integrated assessment of sleep, cognition, and neuropathology following long-term trazodone treatment in a mouse model of AD. In participants with probable AD with sleep disturbances, trazodone for two weeks increased total time spent asleep (avg 43 min), sleep efficiency, and circadian rhythm amplitude^27,33^. Another study, a 4-year retrospective evaluation of the UCSF Memory and Aging cohort, found that those treated with trazodone had slower cognitive decline (MMSE; 2.6-fold faster decline in non-users), and the benefit of trazodone was also seen when looking only at individuals diagnosed with AD^34^. In a recent preclinical study using the APP/PS1 mouse model of AD, oral administration of trazodone for 2 weeks alleviated Aβ pathology and reduced circulating soluble ST2, a potential disease-modifying factor in AD^41^. The authors further showed that trazodone suppresses sST2 expression by antagonizing α1- and β-adrenergic receptor signaling^41^. Similarly, another preclinical study using the rTg4510 tauopathy mouse model showed that trazodone improved memory performance and reduced tau accumulation and neuroinflammation^42^. Together, these findings support the therapeutic potential of trazodone in AD and suggest that trazodone may remain beneficial as pathology progresses, whereas other sleep-targeting medications may exert effects only at earlier stages^22,23^.

In addition to the potential disease modifying effects of SWS promotion, trazodone also acts directly on downstream cellular pathways that offer neuroprotection. Trazodone acts on the unfolded protein response (UPR) pathway, which is overactivated in AD and a promising therapeutic target for protein misfolding neurodegenerative diseases^43^. Additionally, trazodone exerts high binding affinity for multiple serotonin receptors, namely 5-HT2A, α1-adrenergic and SERT receptors. Notable in this regard, is that chronic, independent inhibition of these receptors using selective inhibitors reduce amyloid plaque deposits.^44,45^ Trazodone as a multimodal antagonist against multiple receptors in the serotonin pathway while being a well-tolerated medicine therefore renders itself an attractive disease modifying candidate for slowing amyloid pathologies. By treating APP^NL-F^ mice for 60 days, we were able to observe alleviated neuropathology at vulnerable regions associated with AD. To confirm whether the observed therapeutic outcomes are effects of promoting NREM sleep or downstream mechanisms from blocking serotonin receptors, mouse models with conditional knockout or knockdown of these receptors in the brain may be a useful tool for mechanistic investigation *in vivo*.^46,47^ Furthermore, selective enhancement of NREM sleep via other methods, such as applying optogenetic stimulation to enhance slow oscillation at comparable levels as observed in our trazodone-treated mice,^48^ could be useful to identify the mechanisms underlying the disease modifying effects of trazodone.

In our study, although females treated with trazodone showed sleep consolidation effects and reduced amyloid pathology comparable to the treatment effects in males, the females did not exhibit the lower levels of microglial activation observed in male mice treated with trazodone. Future research should confirm these effects with a larger sample size and investigate the underlying mechanisms to determine whether trazodone’s effects on downstream cellular pathways driving neuroinflammation differs between sexes. It is also worth noting that previous reports showing reduced neuroinflammation by trazodone were conducted in mouse models with profound baseline neuroinflammation and neurodegeneration.^42,43^ In contrast, the APP^NL-F^ model does not exhibit substantial neuroinflammation nor neuronal loss at the investigated ages,^49,50^ hence significant dampening of overall glial activation by trazodone was not expected. For this reason, we also investigated congophilic amyloid plaque-associated microglia, which is one histological feature of neuritic plaques that are highly correlated with the progression of AD.^39,51^ Our findings suggest that chronic trazodone may be effective at lowering microglial activation engaging with neuritic plaques in a sex-dependent manner. This effect did not appear to exacerbate plaque deposition, but instead, could be a result of attenuated congophilic plaque burden, albeit its underlying mechanism remaining unclear. Interestingly, one recent report demonstrated differential labeling specificities of amyloid plaques by X-34 (both fibrillar and dense-cored plaques) and Thioflavin-S (dense-cored plaques) in APP/PS1 mice, which are generally believed to represent the intermediate and late stages of amyloid plaque maturation, respectively.^52^ This is consistent with our findings showing lower Thioflavin-S relative to X-34 labeling in all investigated brain regions. Therefore, the overall downtrend on Thioflavin-S labeled deposits observed in the trazodone-treated mice in the 14-month cohort perhaps indicate slowing of the transition of fibrillar plaques into dense-cored plaques, a process in which microglia are believed to contribute to.^52^ Future research can focus on elucidating potential mechanisms of chronic trazodone treatment modulating microglia in their processing of amyloid plaques.

Finally, chronic trazodone treatment in the APP^NL-F^ mice was also associated with functional benefits, specifically enhanced short- and long-term recognition memory. These beneficial effects of trazodone on recognition memory support previous reports from mouse models^42,43^ as well as the retrospective clinical report of slower cognitive decline in trazodone users.^34^ The functional benefits of trazodone treatment may be driven by the enhancement of slow-wave sleep quality, which could support overnight memory consolidation,^55^ the reduced AD neuropathology in the hippocampus, or other neuroprotective effects.^43,56^

In conclusion, chronic trazodone treatment promotes sleep consolidation through slow wave sleep enhancement, ameliorates amyloid-plaque burden and neuroinflammation, and improves short-term and long-term recognition memory in a mouse model of AD. These findings support the therapeutic potential of repurposing trazodone for AD, which is widely used and well-tolerated by older adults at low, sleep-promoting doses.

## Methods

### Animals

All experimental protocols were approved by the Simon Fraser University Animal Care and Use Committee (Protocol #1365P-23). APP^NL-F^ are knock-in mice that carry the Swedish (KM670/671NL) and Beyreuther/Iberian (I716F) mutations^50^. APP^NL-F^ mice were provided by Cheryl Wellington at the University of British Columbia. Three cohorts of the homozygous APP^NL-F^ mice were used. Two cohorts underwent EEG/EMG recording and neuropathological analysis: a 9-month-old cohort (*n* = 16; 9 trazodone, 7 vehicle control; 7 females across groups) and a 14-month-old cohort (*n* = 19; 11 trazodone, 8 vehicle control; 9 females across groups), representing early and intermediate stages of amyloid pathology, respectively. Some animals were excluded from EEG power analysis due to high artefact in EEG, although all remained suitable for vigilance state scoring, which are supported by the second EEG channel, as well as EMG and video recordings. For the power spectra analyses, one trazodone-treated male and one control male from the 9-month-old cohort, and three control males from the 14-month-old cohort were excluded. Two trazodone-treated females and one trazodone-treated male from the 14-month-old cohort were excluded only from baseline power spectra analyses. For histological analyses: one control female mouse was excluded from glial and tau analyses due to temperature-related damage from the EEG/EMG headcap on the day of tissue extraction. This mouse was included for 6E10 and Thioflavin-S labeled amyloid analyses as we did not find abnormalities on the levels of both soluble and insoluble amyloid associated with this artifact. Mice in these cohorts were singly housed and maintained on a 12:12-h light/dark cycle. A third cohort of 14-month-old mice (*n* = 32; 17 trazodone, 15 vehicle control; 15 females across groups) was used for behavioral testing. One animal was excluded due to stereotypic behavior. Additional animals were excluded as outliers using the 1.5 × interquartile range rule: two trazodone-treated males and one trazodone-treated female for the 1-h delay condition, and two trazodone-treated females for the 24-h delay condition. Mice in this cohort were housed in same-sex pairs, and maintained on 11:13-h light/dark cycle to better accommodate the behavioural testing logistics.

### Trazodone administration

Each cohort was divided into two groups: a trazodone treatment group and vehicle control group. As previously described^36^, the treatment group received palatable food mixed with trazodone (60 mg/kg) and the control group received palatable food mixed with distilled water, daily for 2 months in the EEG/EMG and neuropathology cohorts and for 4 months in the behavioural testing cohort (Fig. 1). The treatments were prepared immediately prior to the daily administration, by dissolving trazodone hydrochloride (HCl) powder (catalog number T6154; Sigma-Aldrich) in sterile water to achieve 10-50mg/ml concentration and then mixed with a palatable food (VitaKraft® Drops with Strawberry, Product #25451, Vitakraft Sunseed, Bowling Green, OH, United States). Voluntary oral administration was necessary because oral gavage cannot be done safely and effectively once the EEG/EMG are implanted. Trazodone is also too acidic to humanely conduct daily injections in mice without the addition of a solvent or suspension-agent, which introduce the potential for unintended physiological effects.

Mice were food restricted to maintain 85-97% of baseline body weight with standard lab chow food pellets provided daily at ZT13 (1h after light-off) for the EEG-neuropath cohorts and ZT15 (4h after lights off) for the behavioural cohort, to time feeding after the daily behavioural testing was complete. Once the body weight was within the target range, the animals were introduced to the palatable food without trazodone daily at ZT 23 (1h before light-on). When animals started to reliably ingest the palatable food within 15 min, the animals undergoing EEG recording were habituated to the EEG recording chamber for one day and then recorded for 24 h to establish baseline EEG. The trazodone treatment was then administered daily. At the beginning of the treatment, the dose was increased gradually to encourage ingestion: 10mg on day 1, 40mg on day 2, and then 60mg day 3 to day 60. If a mouse did not consume the entire dose, it was given a lower dose (10 or 40 mg/kg) the next day to encourage consumption, and then increased back to 60mg/kg. Dosing was adjusted daily based on daily body weight, recorded at ZT23.

### EEG implantation and recordings

Mice were implanted with a 2-channel EEG/1-channel EMG headmount (#8201-SS; Pinnacle Technology, Lawrence, KS, United States) (methods previously described^36,57^). Throughout the EEG implantation procedure, animals were anesthetized deeply with isoflurane. The cranial implant consisted of four stainless steel screws at coordinates relative to bregma: AP: +/− 3 mm, ML: +/− 1.5 mm. The screw electrodes were inserted through holes of a prefabricated EEG headmount and rotated into 4 burr holes drilled through the skull. Two EMG electrode wires attached to the headmount were inserted under the nuchal muscles. A layer of dental cement was applied to secure the headmount to the skull. Meloxicam (5mg/kg, IP), buprenorphine (0.07mg/kg, SC), and lidocaine (7mg/kg, SC along the incision site) were administered for analgesia at the beginning of the surgery. Lactated Ringer’s (10mg/kg) was administered at the end of surgery to prevent dehydration. Meloxicam and buprenorphine were also provided for 2 days post-op. All mice were given at least 7 days for recovery before EEG recordings.

The animals were habituated for at least 24 h to a rectangular clear plexiglass recording cage and the wireless EEG unit (#8274-SL; Pinnacle Technology). After the habituation period, all animals were recorded with an *in vivo* EEG monitoring system (#8200-K1-SE3, 8236; Pinnacle Technology) for one day as a baseline and then 60 days while receiving the treatment (palatable food mixed with trazodone 60mg/kg) or vehicle (palatable food mixed with distilled water). The recording chamber was maintained on a regular 12:12 LD cycle with *ad libitum* access to water. The EEG signals were recorded at a sampling rate of 256 Hz, with a total gain of 2600 V/V. A high-pass filter was set at 0.5 Hz for EEG recordings and 10 Hz for EMG recordings. All animals underwent EEG recording on the baseline day and then in 2-day sessions (1 day habituation, 1 day recording) every 7–10 days throughout the 60-day treatment period.

### Sleep Analysis

For vigilance state and power spectra analyses, EEG traces were scored for wake, NREM sleep, and REM sleep in 10-s epoch durations using Sirenia Sleep Pro software (Pinnacle Technology) by an investigator blind to the treatment condition (methods previously described^15,16,36,57^). Briefly, the data were first cluster scored by grouping epochs based on EEG and EMG frequency bands (e.g., slow oscillation, delta, theta, alpha, beta, gamma), identifying bouts of NREM, REM and wake periods. Each epoch’s classification was then visually confirmed by reviewing EEG and video recordings and corresponding spectral plots. Wake was defined by low-amplitude EEG (dominant frequency above 4 Hz) and high-amplitude EMG. NREM sleep was defined by high-amplitude EEG and frequencies below 4 Hz, and low-amplitude EMG. REM was defined by EEG peak frequencies between 4 and 8 Hz, uniform low amplitude EEG waveforms, low-amplitude EMG, and transitioning from NREM. Epochs were categorized based on the predominant state (>50%) of each epoch. To analyze the power spectrum, frontal EEG data were used. Following the Fast Fourier Transform, a bandpass filter with a range of 0.5–50 Hz was applied to the data. The frequency bands were defined as slow oscillation (0.5–1Hz), delta (1–4 Hz), theta (4–8 Hz), alpha (8–12 Hz), beta (12–30 Hz), and gamma (30–50 Hz).

### Tissue Collection and Processing

Immediately after the final EEG recording, mice were sacrificed by deeply anesthetizing with 150 mg/kg ketamine and 20 mg/kg xylazine, then quickly perfused with ice-cold PBS. The extracted brains were medially bisected. One hemibrain was snap-frozen on dry ice and stored at −80°C until processing, and the other hemibrain was fixed in 4% paraformaldehyde, followed by cryopreservation in 30% sucrose, prior to cryosection into 40 μm sections. Frozen hemibrains were serially homogenized as previously described^58^ to extract Aβ in both carbonic (soluble) and guanidine hydrochloride (GuHCl) (insoluble) fractions. Protein concentrations were determined by Lowry-based protein assay (BioRad).

### Histology

Three coronal brain sections were sampled from −1.5 mm to −2.3 mm, and −2.8 mm to −3.6 mm from bregma for each mouse. Total amyloid plaque burden was analyzed by immunohistochemistry. After pre-treating sections with 88% formic acid for 5 min, brain sections were permeabilized and blocked with 0.25% Triton X-100, 0.3% hydrogen peroxide and 5% nonfat milk, followed by overnight primary antibody incubation (6E10, 1:1000, BioLegend #803004). After washing, sections were incubated with biotinylated secondary antibody (1:1000, Southern Biotech #1071-08), then with ABC substrate (VectorLab #PK-6100) followed by 3,3’-Diaminobenzidine (DAB). Fibrillary-cored amyloid plaque burden was assessed by either 1% Thioflavin-S or 25 µM X-34. Sections were mounted and coverslipped with VECTASHIELD Vibrance Antifade Mounting Medium (Vector Laboratories #H-1700-2). Microglial and astroglial activity were assessed by immunofluorescence. After washing, brain sections were permeabilized and blocked with 5% normal goat serum in 0.25% Triton X-100, then incubated with primary antibodies (Iba1, 1:1000, WAKO #019-19741; GFAP GA5, 1:500, Invitrogen #53-9892-82). Brain sections were then washed and incubated with Alexa Fluor 594 conjugated anti-rabbit secondary antibody (Invitrogen, #A11012). To assess microglia surrounding fibrillary-cored plaques, brain sections were first stained with X-34 dye, followed by immunostaining with anti-Iba1 antibody. All slides were imaged on the Axio Scan Z1 at 20x magnification and tiled to generate the full section image using Zen software (Zeiss).

### Enzyme-linked Immunosorbent Assay (ELISA)

Carbonic and GuHCl-buffered hemibrain lysate were used to quantify soluble and insoluble Aβ_40_and Aβ_42_ using human Aβ_40_ (Invitrogen #KHB3481) and Aβ_42_ (Invitrogen #KHB3441) ELISA kits according to manufacturer’s instructions. Carbonic and GuHCl-buffered brain lysates were diluted to reach detection sensitivity of the kit. Absorbance values were obtained on the Synergy Neo2 plate reader (BioTek). Each sample was assayed in triplicate, and the average absorbance value of each triplicate was reported.

### Immunoblot

The carbonic fraction of hemibrain lysate was analyzed for total- and phospho-tau by DA9 and PHF1 antibodies (generous gift from Dr. Peter Davies), respectively. 25 µg of lysate was resolved in 4-15% Mini-PROTEAN precast gel (Bio-Rad #4561086), followed by transferring onto nitrocellulose membrane. After blocking with 5% BSA, the membrane was incubated with primary antibodies overnight. After washing and developing with secondary antibody, the membrane was imaged on the ChemiDoc Imaging System (Bio-Rad). β-actin was used as a housekeeping gene.

### Novel object recognition

Novel object recognition testing began after approximately 130 days of trazodone treatment. Mice were habituated to the maze for 5 min per day on three consecutive days. The maze was a white, opaque Y-maze containing three identical arms and two sliding platforms for object placement. During the sample phase, mice were allowed to explore two identical objects for 5 min. Following a delay of either 1 or 24 h, mice underwent a 5-min test phase in which one familiar object was replaced with a novel object. Object pairs were similar in size and were validated before testing to exclude biased preferences. Object identities and locations were counterbalanced across sex and treatment groups. Testing was conducted during the active phase (ZT11-ZT15) under dim light. The discrimination index (DI) was calculated as the time spent exploring the novel object divided by the total time spent exploring both objects during the test phase^40^. Additional procedural, scoring, and exclusion details are provided in the Supplementary Methods.

### Data analysis

Power spectra were analyzed as a percentage of total power across the frequency range of 0.5 Hz to 50 Hz. Statistical analyses were conducted using GraphPad Prism (version 10.5.0). All sleep analyses (Fig. 2,3) were performed using two-way ANOVA, or mixed-effects models when there were missing values, followed by Tukey’s or Šidák’s multiple comparisons tests when a significant interaction effect was detected.^73,74^ All histology image analyses were performed on ImageJ (NIH). The regions of interest (ROI) were defined according to anatomical landmarks referenced from the Allen Mouse Brain Atlas (https://mouse.brain-map.org/experiment/thumbnails/100048576?image_type=atlas) to isolate temporal cortex (TCTX), hippocampus (HPC), and entorhinal cortex (ENT). Total and fibrillary-cored amyloid plaque burden, and total microglia were analyzed and reported as percentage area within ROI. Analysis of microglia surrounding congophilic amyloid plaques was performed by quantifying the percentage area of immunoreactive cells within a circular region at a radius of 50 μm projected from the circumference of X-34-labeled plaques within each ROI (Extended Data Fig. 6). This distance was selected according to a previous report showing increased densities of microglia proximal to Thioflavin-S-labeled plaques in post-mortem human brain sections with AD.^39^ Histological and biochemical data were analyzed by two-way ANOVA, followed by Bonferroni multiple comparisons tests. Densitometry of immunoblot analysis was conducted using ImageLab software (Bio-Rad) and analyzed by student-t. For novel object recognition, discrimination indices for the 1- and 24-h retention intervals were analyzed separately using two-way ANOVAs, with treatment and sex as factors. All data are presented as mean ± standard error of the mean (SEM), and all tests were two-tailed. Complete statistical results are provided in Supplementary Table 1.

## Acknowledgements

We would like to thank the SFU animal research staff for their support and care for the animals used in this study. We also thank the following research assistants: Erin Asu, Dana Braynina, Seoyoung Chae, Mona Chen, Artemis Kohanfekr, Katherine Mantel, Sarah McGuire, Dulni Peramunugamage, Daniela Purvica, Gisany Ravichandran, Afnan Sahibzada, Kiana Shakiba, Sofiya Soboleva, and Elizabeth Soulliere.

## Author Contributions

M.A: investigation, methodology, project administration, visualization, and writing - original draft. J.Y.: investigation, methodology, visualization, writing - review & editing. E.S.: investigation, visualization, writing - review & editing. C.J.S: investigation. H.H: investigation. R. G.: investigation, project administration. T. Y.: investigation. H.H. F: writing - review & editing. C.L.W: resources, writing - review & editing, B.K.: conceptualization, methodology, resources, supervision, funding acquisition. All authors reviewed and approved the final manuscript.

## Preprint Information

This manuscript has been submitted to bioRxiv; the DOI is pending.

---

**Extended Data Fig. 1.**
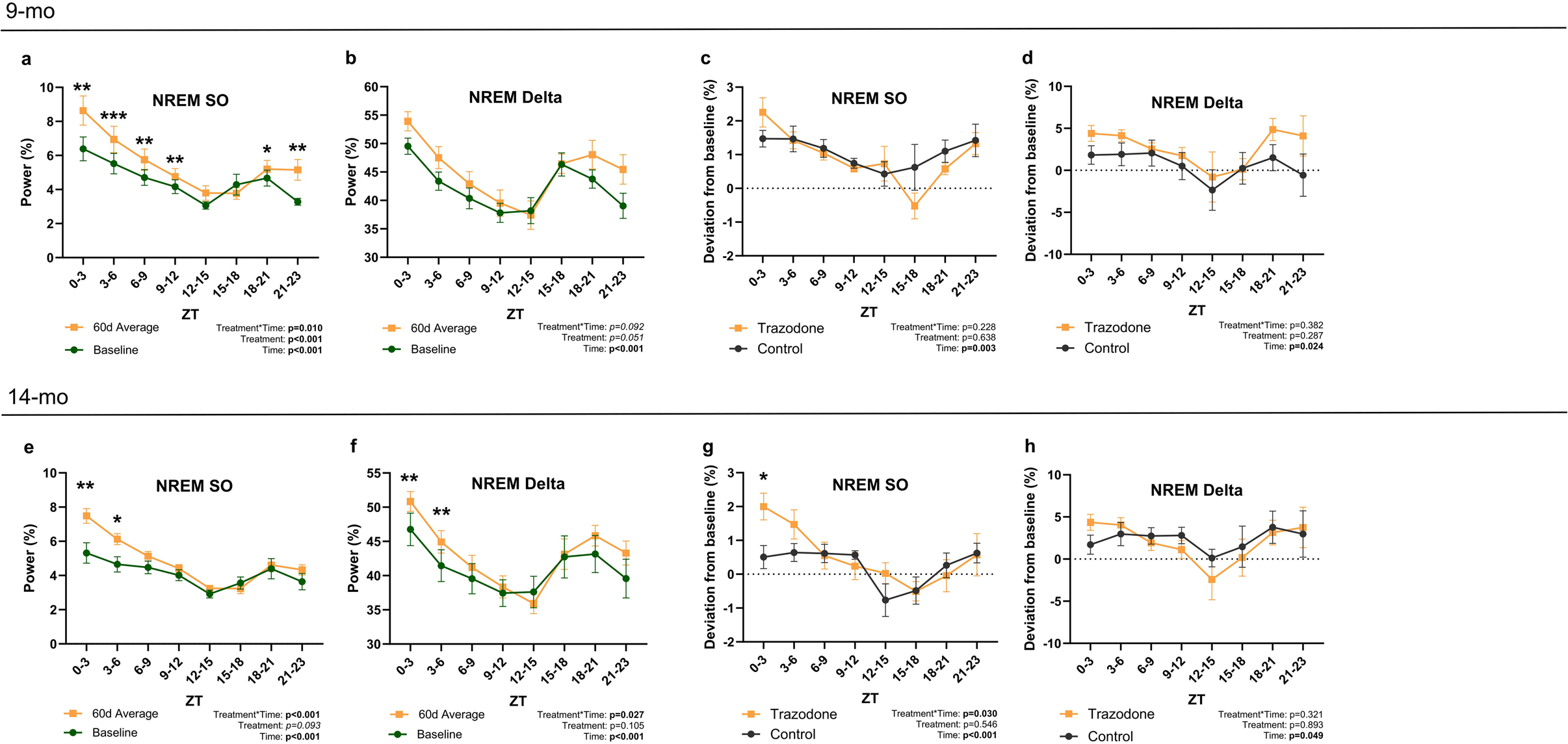
Trazodone increases NREM SO and delta power at the beginning of the 12-hour light phase. Effects of chronic trazodone treatment on NREM EEG power during ZT0-23 in APP^NL-F^ mice treated daily from 9 to 11 months of age (a–d) and from 14 to 16 months of age (e–h). **(a–d)** 9-month cohort: NREM SO and delta power in 3-h bins, comparing baseline with the 60-day treatment average (a, b) or vehicle- and trazodone-treated APP^NL-F^ mice across the day (c, d). **(e–h)** 14-month cohort: NREM SO and delta power in 3-h bins, comparing baseline with the 60-day treatment average (e, f) or vehicle- and trazodone-treated APP^NL-F^ mice across the day (g, h). Data (mean ± SEM) were analyzed by two-way ANOVA followed by Tukey’s multiple comparisons test, or mixed-effects models when there were missing values, followed by Šidák’s multiple comparisons tests. * p<0.05, ** p<0.01, *** p<0.001.

**Extended Data Fig. 2.**
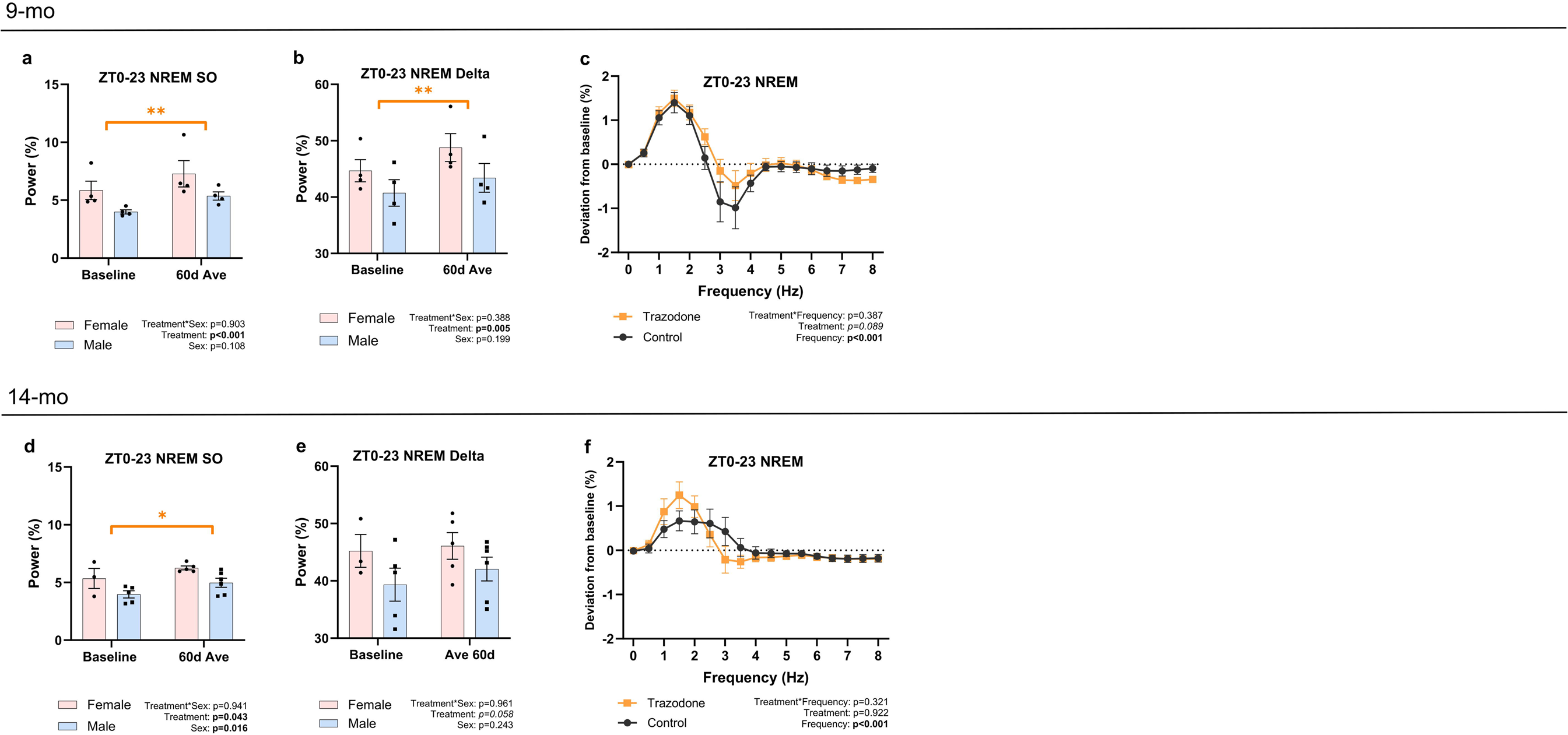
Trazodone increases NREM SO and delta power during ZT0–23 similarly in both sexes (a, b, d &. **e)** Comparison across sexes of ZT0–23 NREM SO and delta power at baseline and averaged across the 60-day treatment period. Sex differences were present at baseline, but the effects of trazodone did not differ between sexes**. (c & f)** ZT0–23 NREM power deviation from baseline in vehicle-treated control versus trazodone groups (60-day average). Data (mean ± SEM) were analyzed by two-way ANOVA followed by Tukey’s multiple comparisons test, or mixed-effects models when there were missing values, followed by Šidák’s multiple comparisons tests. * p<0.05, ** p<0.01, *** p<0.001.

**Extended Data Fig. 3.**
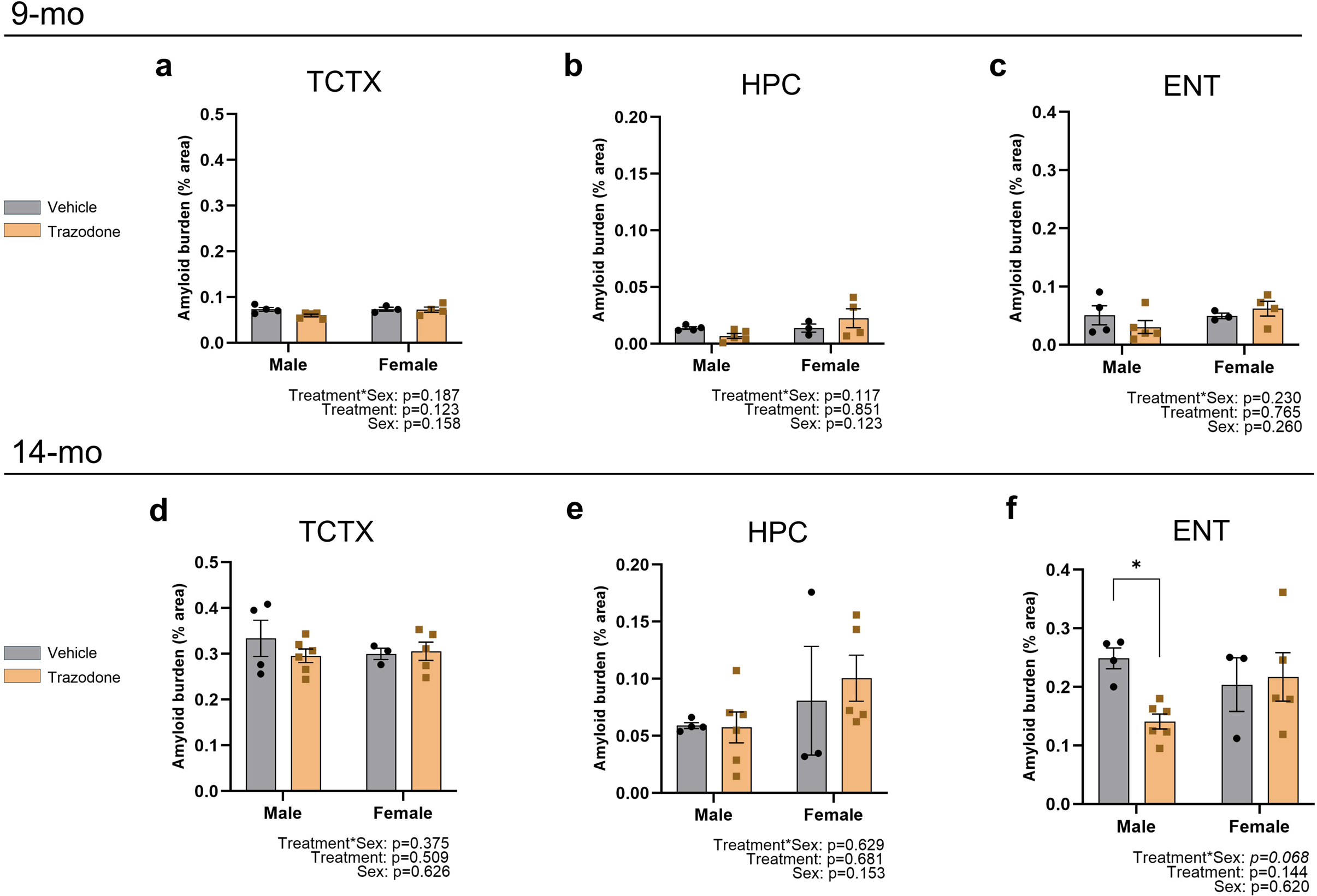
Trazodone reduces X-34 labeled congophilic amyloid plaque in the entorhinal cortex of male mice. Effects of chronic trazodone treatment on X-34 labelled fibrillary-cored amyloid plaques in the brain regions of APP^NL-F^ mice treated daily from (a–c) 9 to11 month of age and (d–f) 14 to 16 month of age. Quantification of X-34 labeled amyloid plaque burden within **(a, d)** temporal cortex, **(b, e)** hippocampus, and **(c, f)** entorhinal cortex. The percentage area of X-34 pixels within each brain region are reported in the bar graph. Each data point represents one sample. Data (mean ± SEM) were analyzed by two-way ANOVA followed by Bonferroni multiple comparison post-hoc test. * p<0.05.

**Extended Data Fig. 4.**
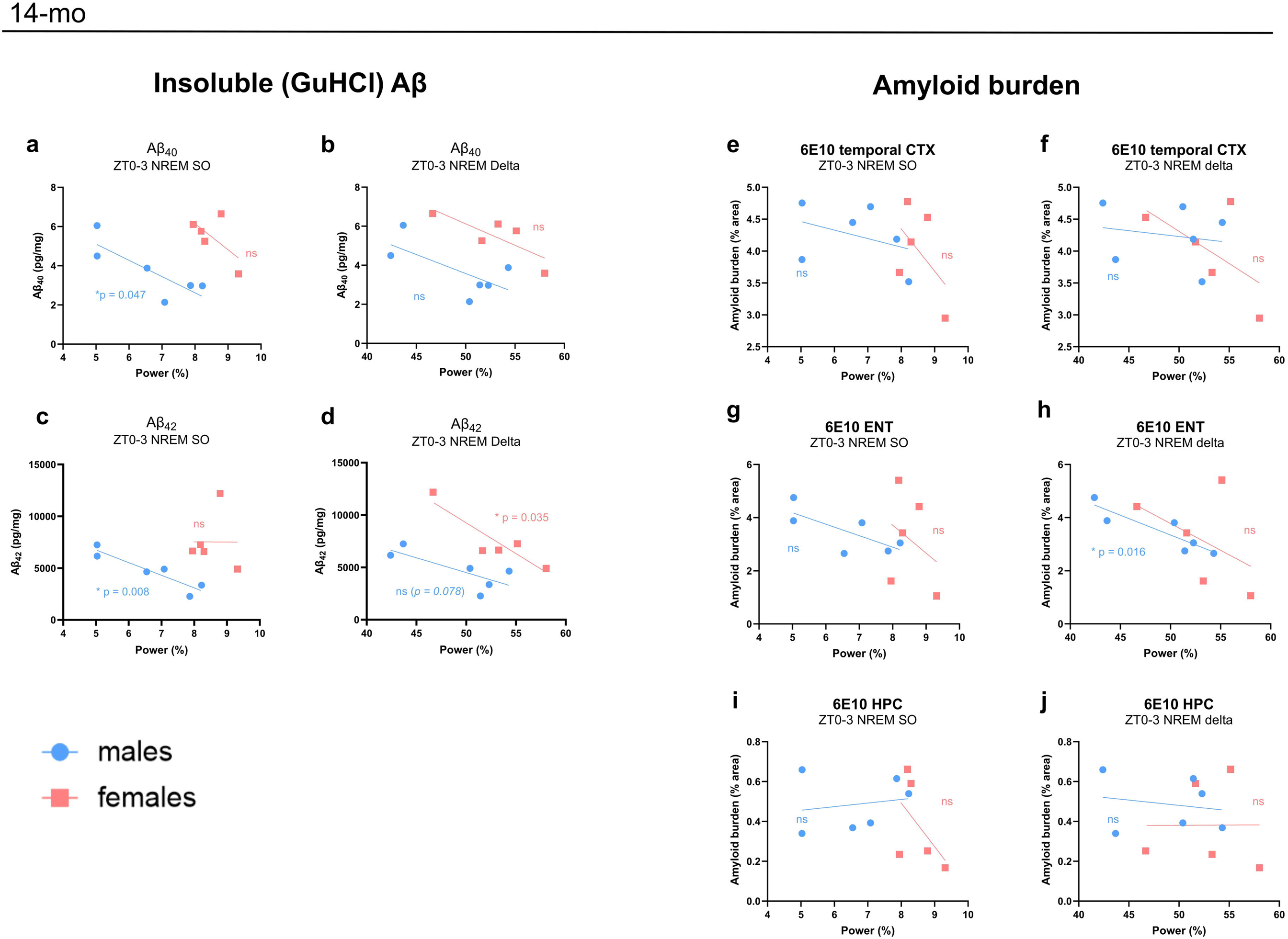
NREM slow-wave power is associated with levels of neuropathology. Correlation between ZT0–3 NREM SO and delta power and neuropathological measures in APP^NL-F^ mice treated daily with trazodone from 14 to 16 months of age. **(a– d)** Simple linear regression of NREM power (SO or delta) versus insoluble Aβ quantified by ELISA: (a, b) Aβ40 and (c, d) Aβ42. **(e–j)** Simple linear regression of NREM power (SO or delta) versus amyloid burden (% area labeled by 6E10): (e, f) temporal cortex, (g, h) entorhinal cortex, and (i, j) dorsal hippocampus. * p<0.05.

**Extended Data Fig. 5.**
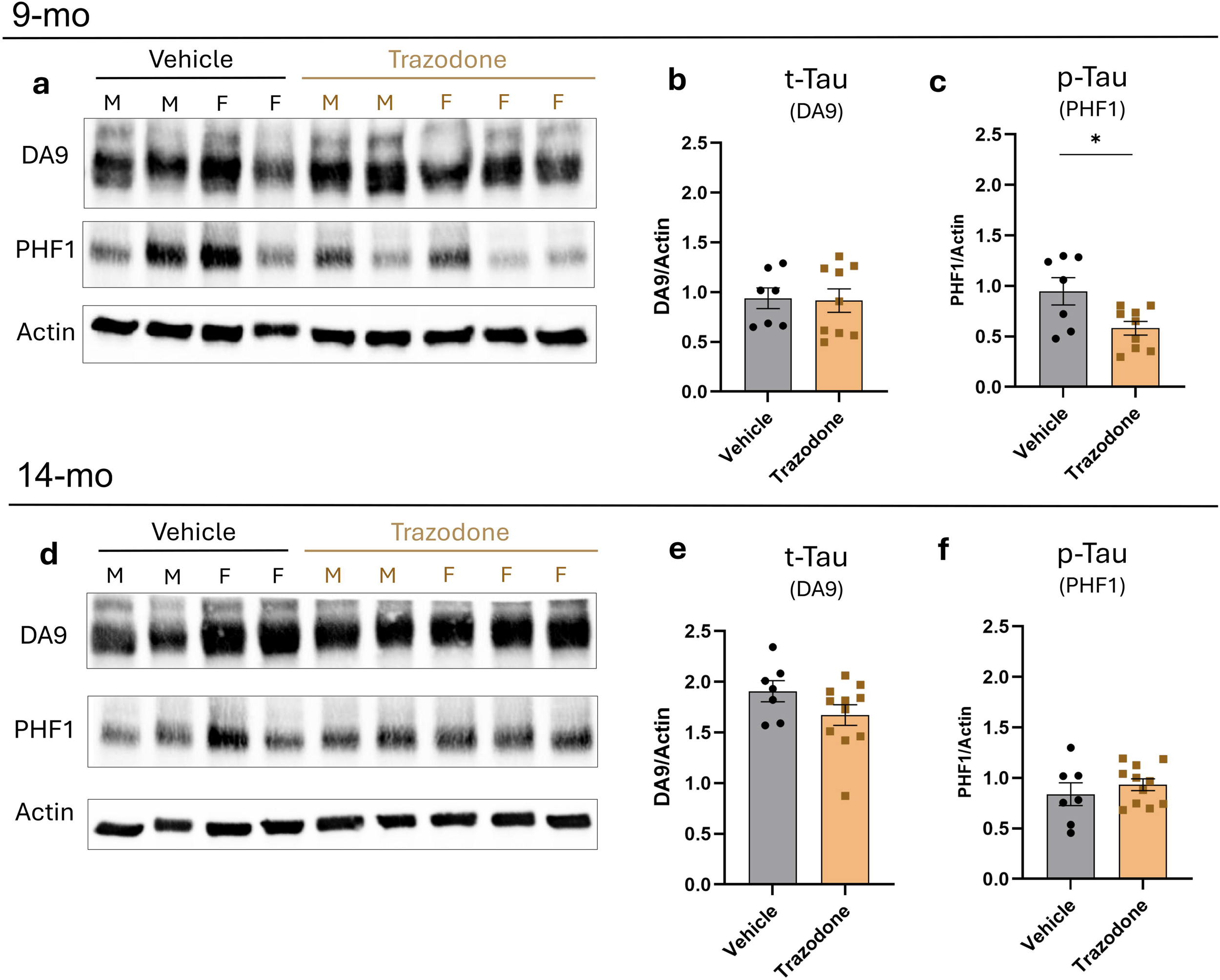
Trazodone alters tau phosphorylation in the brain. Effects of chronic trazodone treatment on the levels of total (t-Tau) and phosphorylated tau (p-Tau) in APP^NL-F^ mice treated daily from 9 to 11 months of age (a–c) and from 14 to 16 months of age (d–f). Representative blot images of total tau detected by antibody DA9 and PHF1 (a, d). Densitometry of total tau (b, e) and phosphorylated tau (c, f). Actin is used as control for normalization. Data (mean ± SEM) were analyzed by student-t. *<0.05.

**Extended Data Fig. 6.**
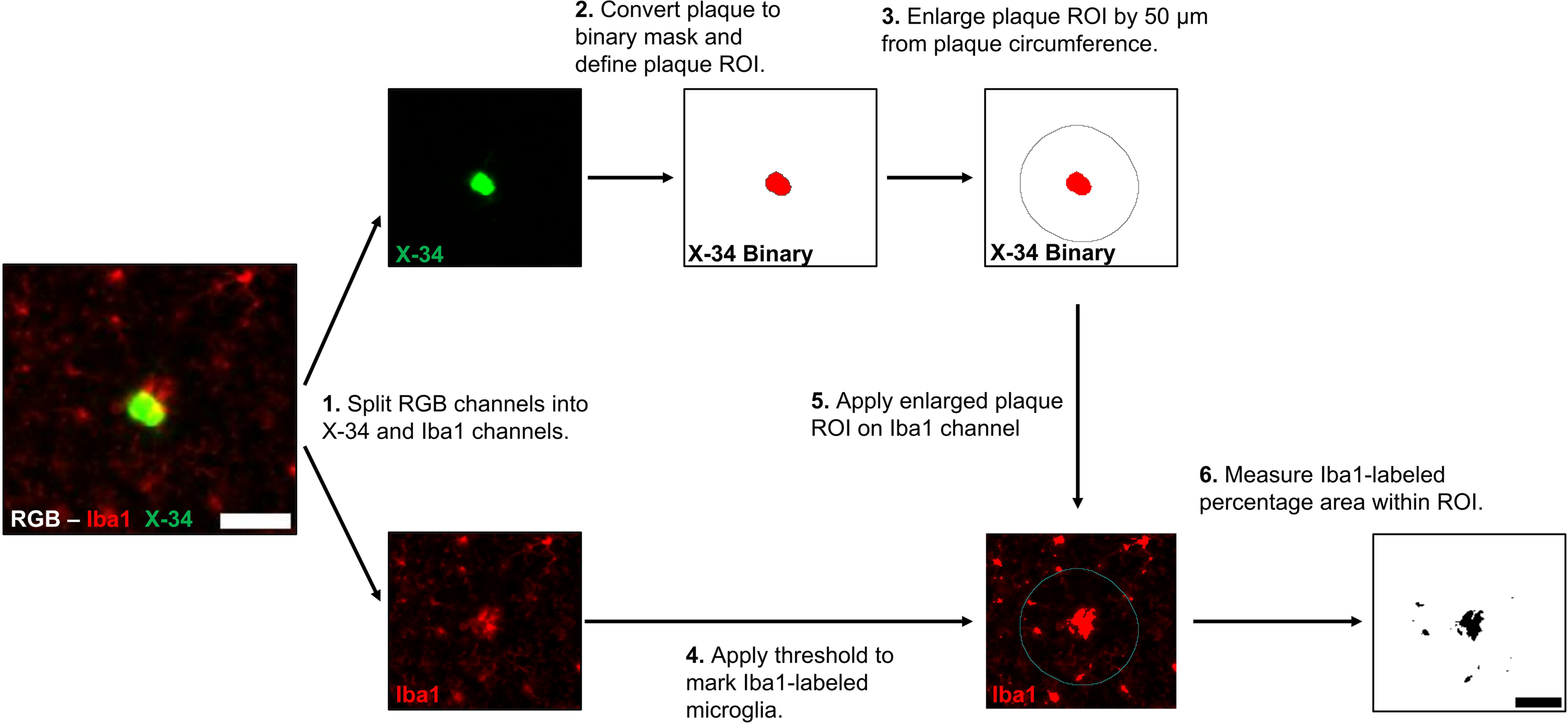
Summary of analysis of plaque-associated microglia on ImageJ. A flow chart describing the process in measuring microglial activity interacting with congophilic amyloid plaque. 1^st^: RGB images are split into red (Iba1) and X-34 (green). 2^nd^: Thresholding is applied to mark X-34 labeled plaques, then converting to binary mask to enable marking the plaque circumference as new ROI (black line tracing plaque). 3^rd^: Enlarge plaque ROI by projecting 50 µm outward (black circle). 4^th^: Apply enlarged plaque ROI (cyan) onto Iba1 channel. 5^th^: Apply thresholding to label microglia. 6^th^: Perform “Analyze particles” on ImageJ to measure Iba1-positive percentage area within enlarged plaque ROI. A macro pipeline was generated to analyze plaque-associated microglia within brain regions.

