## Supplementary Methods for "Chronic trazodone treatment consolidates sleep, improves memory, and reduces amyloid pathology in a mouse model of Alzheimer’s disease"

### Behavioural Cohort

Prior to object recognition testing, the mice in the behavioural cohort underwent touchscreen cognitive testing not described here (R.G., T.Y., H.H., M.A., J.Y. and B.A.K., in preparation). The 11:13 LD schedule was implemented to extend the time between the food rewards used during touchscreen testing and the treatment, which was administered with a palatable food.

### Novel Object Recognition

An overhead camera was used to record the entire Y-maze. Testing times were counterbalanced across sex and treatment groups. During novel object recognition testing, objects were secured to the sliding platforms with Blu Tack, which was replaced between phases. Testing was completed approximately 15 min before feeding. The maze and objects were cleaned with 70% ethanol between mice, and at least a 72-h interval was imposed between trials to maintain exploratory motivation and minimize interference from previous trials.

Exploration was defined as orienting the head toward and sniffing an object from a distance of ≤5 cm. Looking at an object without approaching it, remaining near its base, sitting or climbing on it, and making physical contact without active investigation were not scored as exploration. Videos were scored using JWatcher v1.0, with the scorer blinded to treatment and object novelty.

Trials were excluded if total object exploration was <3 s during either the sample or test phase[59]). Outliers were identified using the 1.5 × interquartile range rule. Trials in which a mouse overturned an object and gained access to the mounting putty were also excluded.
